# Guidelines for Evaluating the Comparability of Down-Sampled GWAS Summary Statistics

**DOI:** 10.1101/2023.03.21.533641

**Authors:** Camille M. Williams, Holly Poore, Peter T. Tanksley, Hyeokmoon Kweon, Natasia S. Courchesne-Krak, Diego Londono-Correa, Travis T. Mallard, Peter Barr, Philipp D. Koellinger, Irwin D. Waldman, Sandra Sanchez-Roige, K. Paige Harden, Abraham A Palmer, Danielle M. Dick, Richard Karlsson Linnér

**Affiliations:** Department of Psychology and Population Research Center, University of Texas at Austin; Department of Psychiatry, Robert Wood Johnson Medical School, Rutgers University; Population Research Center, the University of Texas at Austin; Department of Economics, School of Business and Economics, Vrije Universiteit Amsterdam; Department of Psychiatry, University of California San Diego; Psychiatric and Neurodevelopmental Genetics Unit, Center for Genomic Medicine, Massachusetts General Hospital, Boston, MA, USA; Department of Psychiatry, Harvard Medical School, Boston, MA, USA; Stanley Center for Psychiatric Research, Broad Institute of MIT and Harvard, Boston, MA, USA; Department of Psychiatry and Behavioral Sciences, SUNY Downstate Health Sciences University; Department of Economics, Vrije Universiteit Amsterdam; Department of Psychology, Emory University; Department of Psychiatry, University of California San Diego, La Jolla, CA, USA; Department of Medicine, Division of Genetic Medicine, Vanderbilt University, Nashville, TN, USA; Institute for Genomic Medicine, University of California San Diego; Rutgers Addiction Research Center in the Brain Health Institute, Department of Psychiatry, Robert Wood Johnson Medical School, Rutgers University; Department of Economics, Universiteit Leiden

**Keywords:** Genomic SEM, summary statistics, data removal, down-sample, leave-one-out, meta-analysis, genomics, genome-wide association study

## Abstract

Proprietary genetic datasets are valuable for boosting the statistical power of genome-wide association studies (GWASs), but their use can restrict investigators from publicly sharing the resulting summary statistics. Although researchers can resort to sharing down-sampled versions that exclude restricted data, down-sampling reduces power and might change the genetic etiology of the phenotype being studied. These problems are further complicated when using multivariate GWAS methods, such as genomic structural equation modeling (Genomic SEM), that model genetic correlations across multiple traits. Here, we propose a systematic approach to assess the comparability of GWAS summary statistics that include versus exclude restricted data. Illustrating this approach with a multivariate GWAS of an externalizing factor, we assessed the impact of down-sampling on (1) the strength of the genetic signal in univariate GWASs, (2) the factor loadings and model fit in multivariate Genomic SEM, (3) the strength of the genetic signal at the factor level, (4) insights from gene-property analyses, (5) the pattern of genetic correlations with other traits, and (6) polygenic score analyses in independent samples. For the externalizing GWAS, down-sampling resulted in a loss of genetic signal and fewer genome-wide significant loci, while the factor loadings and model fit, gene-property analyses, genetic correlations, and polygenic score analyses are robust. Given the importance of data sharing for the advancement of open science, we recommend that investigators who share down-sampled summary statistics report these analyses as accompanying documentation to support other researchers’ use of the summary statistics.

## Introduction

The success of genome-wide association studies (GWASs) depends on sample size (Abdellaoui et al., 2023). Accordingly, genetics researchers increasingly depend on public-private partnerships that pool data collected by academic researchers, national biobanks, and private companies. For example, the company 23andMe Inc. contributed an astonishing 2.5 million observations to a recent GWAS of height (Yengo et al., 2022). However, to protect their interests, private companies place restrictions on the public sharing of GWAS summary statistics and require a potentially lengthy and burdensome application process for researchers to gain access. In some cases, researchers’ institutions are unwilling to agree to the legal terms set by private companies in their material transfer agreements. These restrictions pose a challenge to scientific transparency and slow the pace of genetic discovery.

To address this challenge, researchers can publicly share down-sampled GWAS summary statistics that exclude restricted data (Coleman et al., 2020; Lee et al., 2018; Yengo et al., 2022). This is an imperfect solution, as leaving out a large part of the study sample not only reduces power but can also change the genetic etiology of the trait being studied, potentially leading to substantial differences in downstream analyses (de Vlaming et al., 2017). For instance, down-sampling could influence estimates of genetic correlations with other traits, associations in polygenic score analyses, and insights from bioannotation analyses. We are only aware of one study investigating the effects of excluding restricted data from a univariate depression GWAS (Coleman et al., 2020), prior to including them in a meta-analysis of mood disorders. The authors examined the robustness of SNP heritability estimates, genetic correlations, and gene identification. Although they identified fewer variants in the down-sampled analyses, results were otherwise similar, suggesting that excluding data in their study did not markedly change the genetic etiology of their focal phenotype. However, most of the studies providing down-sampled summary statistics have not evaluated the comparability with restricted data counterparts (Lee et al., 2018; Liu et al., 2019; Wray et al., 2018).

There have been few, if any, systematic investigations of how down-sampling affects results from multivariate GWASs. Multivariate GWAS methods, such as genomic structural equation modeling (Genomic SEM; Grotzinger et al., 2019), have become increasingly popular, as there is substantial genetic overlap across psychiatric and behavioral phenotypes. Genomic SEM models the shared genetic architecture among traits with latent factors representing cross-cutting genetic liabilities. Rather than just examining genetic associations with individual phenotypes, Genomic SEM enables the identification of shared genes. As in phenotypic factor analysis, the construct represented by a latent factor could be sensitive to the choice of indicator phenotypes used in the factor analysis, or the construct might be fairly robust to this decision (Johnson et al., 2004, 2008). Using down-sampled univariate GWAS summary statistics as inputs in Genomic SEM could, therefore, identify a genetic factor structure that occupies a different position in genetic multivariate space. Yet, no studies to our knowledge have examined how down-sampling affects multivariate GWAS in the context of Genomic SEM.

Here, we present a systematic approach to assess the comparability of down-sampled summary statistics with their full data counterparts and examine their suitability for typical follow-up analyses. We used externalizing, a latent factor representing a cross-cutting liability to behaviors and disorders characterized by problems with self-regulation, as our model phenotype. A previous multivariate GWAS by the Externalizing Consortium identified several hundred genomic loci associated with an externalizing (EXT) factor, reflecting shared genetic liability among seven indicator phenotypes (Karlsson Linnér et al., 2021): (1) attention-deficit/hyperactivity disorder (ADHD; Demontis et al., 2019), (2) problematic alcohol use (ALCP; Sanchez-Roige et al., 2019), (3) lifetime cannabis use (CANN; Pasman et al., 2018), (4) reverse-coded age at first sexual intercourse (FSEX; Karlsson Linnér et al., 2019), (5) number of sexual partners (NSEX; Karlsson Linnér et al., 2019), (6) general risk tolerance (RISK; Karlsson Linnér et al., 2019), and (7) lifetime smoking initiation (SMOK; Liu et al., 2019). However, the univariate GWASs on two of the seven phenotypes, SMOK and CANN, contain restricted data, which limits public sharing of the summary statistics from this multivariate GWAS (hereafter, the original study).

Therefore, we developed the following six steps to investigate the robustness of down-sampling and applied them to our scenario of assessing the impact of excluding restricted data from the original study (Karlsson Linnér et al., 2021). As an initial check, we suggest testing whether the genetic correlation between full and down-sampled GWASs on the same trait is less than unity, which would suggest imperfectly overlapping genetic etiology. The greater the discrepancy between the genetic correlation of the full and down-sampled GWASs on the same trait, the more important it is to evaluate the comparability of down-sampled analyses.

We recommend that investigators sharing down-sampled GWAS summary statistics report these analyses as documentation for use by other researchers:

1. What is the loss of genetic signal in down-sampled univariate GWASs (which may later be used as indicator phenotypes in Genomic SEM)?
2. How do the factor loadings and factor model fit differ in multivariate Genomic SEM when the indicator phenotypes are down-sampled univariate GWASs?
3. What is the loss of genetic signal at the factor level of multivariate GWAS when the indicator phenotypes are down-sampled univariate GWASs?
4. How similar are gene-property analyses when using down-sampled GWASs?
5. How similar is the pattern of genetic correlations with other traits when using down-sampled GWASs?
6. How much explanatory power is lost when using polygenic scores (PGSs) constructed from down-sampled GWASs?

## Methods

The code for the following analyses is publicly available here: https://github.com/Camzcamz/EXTminus23andMe and the externalizing minus 23andMe summary statistics are available here: https://externalizing.rutgers.edu/request-data/.

### 1. What is the loss of genetic signal in down-sampled univariate GWASs?

The following five key indicators are useful for evaluating the loss of genetic signal in down-sampled univariate GWASs: (1) effective sample size (*EffN*), (2) heritability, (3) mean χ^2^, (4) genomic inflation factor, and (5) attenuation/stratification bias ratio of LD Score regression (see formula in Table 1). *EffN* is a transformation relevant for GWAS on binary traits that transforms an unbalanced number of cases and controls to effectively reflect the sample size of a balanced analysis (i.e., 50% cases). For a meta-analysis of *k* cohort-level univariate summary statistics, it is the sum of *EffN_k_* = 4*V_k_ (1-V_k_)N_k_, where *V_k_* is the cohort-specific proportion of cases, and *N_k_* is the cohort-specific total number of cases and controls. For GWAS on continuous traits, *EffN* can be replaced by the total sample size (*N*). The remaining four key indicators are standard estimates of LD Score regression (version 1.0.1; Bulik-Sullivan et al., 2015).

**Table 1.** Summary of GWAS summary statistics with and without 23andMe data for seven externalizing-related disorders and behaviors

|  | EXT (Karlsson Linnér et al., 2021) |  |  |  |  |  | Down-Sampled EXT (minus 23andMe) |  |  |  |  |  |
| --- | --- | --- | --- | --- | --- | --- | --- | --- | --- | --- | --- | --- |
| *Phenotype* | *Max N*<br>*(EffN)* | *h<sup>2</sup>*<br>*(SE)* | *λ<sub>GC</sub>* | *Mean χ<sup>2</sup>* | *Intercept* | *Ratio* | *Max N*<br>*(EffN)* | *h<sup>2</sup>*<br>*(SE)* | *λ<sub>GC</sub>* | *Mean χ<sup>2</sup>* | *Intercept* | *Ratio* |
| ADHD | 53,293<br>(49,017) | 0.235<br>(0.015) | 1.25 | 1.297 | 1.034 | 0.113 | 53,293<br>(49,017) | 0.260<br>(0.017) | 1.253 | 1.297 | 1.034 | 0.113 |
| ALCP | 164,684<br>(150,640) | 0.055<br>(0.004) | 1.15 | 1.174 | 1.013 | 0.073 | 164,684<br>(150,640) | 0.055<br>(0.004) | 1.149 | 1.174 | 1.013 | 0.073 |
| CANN | 186,875<br>(179,534) | 0.066<br>(0.004) | 1.23 | 1.267 | 1.026 | 0.098 | 164,192<br>(157,230) | 0.068<br>(0.004) | 1.217 | 1.245 | 1.028 | 0.113 |
| FSEX* | 357,187 | 0.115<br>(0.004) | 1.62 | 1.869 | 1.036 | 0.041 | 357,187 | 0.115<br>(0.004) | 1.626 | 1.868 | 1.036 | 0.041 |
| NSEX | 336,121 | 0.097<br>(0.004) | 1.49 | 1.682 | 1.027 | 0.041 | 336,121 | 0.099<br>(0.004) | 1.493 | 1.674 | 1.027 | 0.041 |
| RISK | 426,379 | 0.053<br>(0.002) | 1.37 | 1.461 | 1.019 | 0.041 | 426,379 | 0.053<br>(0.002) | 1.372 | 1.461 | 1.019 | 0.041 |
| SMOK | 1,251,809<br>(1,232,397) | 0.078<br>(0.002) | 2.33 | 3.152 | 1.126 | 0.058 | 652,520<br>(652,518) | 0.079<br>(0.003) | 1.726 | 2.062 | 1.037 | 0.035 |

EXT: Externalizing. Highlighted cells indicate down-sampled summary statistics. *EffN* = sum of cohort-level effective sample sizes. The statistics reported in this table were all estimated with LD Score regression (v1.0.1)(Bulik-Sullivan et al., 2015) : Heritability ( $h^2$ ) is on the observed scale. The genomic inflation factor, $\lambda_{GC}$ , is the median $\chi^2$ statistic divided by the expected median of the $\chi^2$ distribution with 1 degree of freedom. Mean $\chi^2$ is the average $\chi^2$ statistic. Intercept is the estimated LD Score regression intercept. The ratio measures stratification bias, defined as $(\text{intercept} - 1)/(\text{mean } \chi^2 - 1)$ . Abbreviations: ADHD = attention-deficit/hyperactivity disorder; ALCP = problematic alcohol use; CANN = lifetime cannabis use; FSEX = age at first sexual intercourse (reverse coded\*); NSEX = number of sexual partners; RISK = risk tolerance; SMOK = lifetime tobacco initiation. \*Age at first sex was reverse coded so as to expect a positive relationship with EXT.

We down-sampled the univariate GWASs of SMOK and CANN by mirroring the meta-analysis protocol of the original study (Karlsson Linnér et al., 2021) and excluding restricted 23andMe data. We then used these five key indicators to assess the loss of genetic signal in the down-sampled univariate GWASs (Table 1). Finally, we estimated genetic correlations among the seven indicator phenotypes in the down-sampled analysis using LD Score regression (Bulik-Sullivan et al., 2015) and compared them to genetic correlations among the indicator phenotypes in the original study (Figure 1, Table S1).

**Fig 1.**
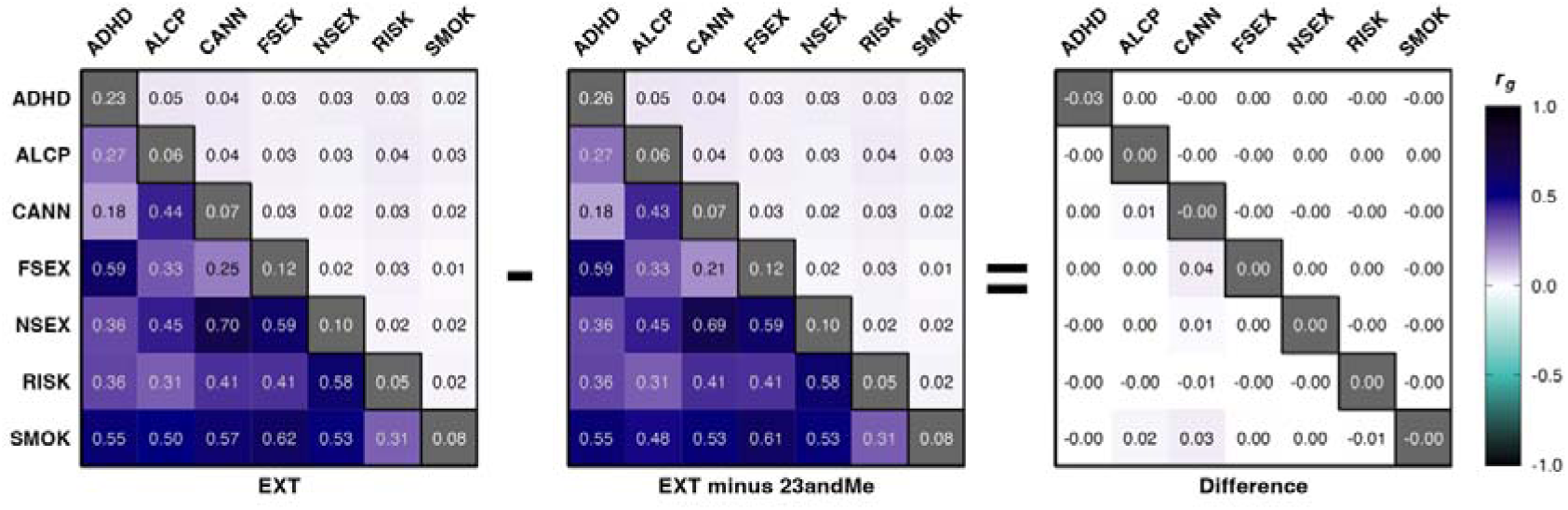
LD Score genetic correlations and heritability estimates for the seven indicator phenotypes of the single-factor models of EXT and EXT-minus-23andMe (see **Step 1**). The left panel displays the analysis of the original study with 23andMe data, the middle panel displays the down-sampled analysis excluding 23andMe data, and the right panel displays the difference in estimates computed by subtracting the values in the middle panel from those in the left panel. The lower and upper triangles display pairwise genetic correlation (*r_g_*) estimates and standard errors, respectively. The diagonals display the observed-scale heritability (*h^2^*; see **Table 1** for standard errors). These results are also reported in **Table S1**. Abbreviations: ADHD = attention-deficit/hyperactivity disorder; ALCP = problematic alcohol use; CANN = lifetime cannabis use; FSEX = age at first sexual intercourse (reverse coded); NSEX = number of sexual partners; RISK = risk tolerance; SMOK = lifetime tobacco initiation.

Stable heritability estimates and attenuation ratios across the original and down-sampled indicators should yield comparable factor loadings in the down-sampled Genomic SEM factor analysis (Step 2), whereas loss of genetic signal, indicated by a decrease in mean χ^2^, should yield larger standard errors in the factor analysis and loss of statistical power to detect SNP effects in the multivariate GWAS (Step 3).

### 2. How do the factor loadings and factor model fit differ in Genomic SEM when the indicator phenotypes are down-sampled univariate GWASs?

Genomic SEM is a flexible modeling approach that (1) estimates an empirical genetic covariance matrix and sampling covariance matrix from input GWAS summary statistics, and (2) evaluates a set of conventional parameters for structural equation modeling, such as factor loadings and residual variances, to minimize the discrepancy between the model-implied and empirical genetic covariance matrices (Grotzinger et al., 2019). Typically, a number of alternative models are compared (e.g., a single-factor model versus a two-factor model) followed by multivariate GWAS to estimate SNP effects on each of the factors in the preferred factor solution (Step 3).

To assess the impact of down-sampling on the factor loadings and model fit, we suggest forcing the best-fitting factor solution from the Genomic SEM analysis of the full dataset (that includes restricted data) onto the empirical genetic covariance matrix of the down-sampled summary statistics, and then evaluating the stability of the factor loadings and factor model fit indicators (e.g., the comparative fit index or the root mean square residual). We do not suggest searching for a better factor solution with the down-sampled indicators because the aim is to evaluate whether down-sampled analyses are representative of their corresponding versions with restricted data.

Thus, we ran the best-fitting Genomic SEM factor model of the original study (Karlsson Linnér et al., 2021): a single-factor model with seven indicator phenotypes (ADHD, ALCP, CANN, FSEX, NSEX, RISK, and SMOK), using unit variance identification of the factor model without SNP effects. However, in the analysis reported here, the input summary statistics for SMOK and CANN were replaced by down-sampled versions (see Step 1). We refer to the original factor model based on analyses with 23andMe data as the EXT factor and the down-sampled version as the EXT-minus-23andMe factor (Table S2).

### 3. What is the loss of genetic signal at the factor level of down-sampled multivariate GWAS when the indicator phenotypes are down-sampled univariate GWASs?

After conducting a multivariate GWAS on the latent factors in down-sampled analyses with Genomic SEM, the loss of genetic signal at the factor level can be assessed by (i) examining the genetic correlation between the respective latent factors of the full and down-sampled summary statistics using bivariate LD Score regression (Bulik-Sullivan et al., 2015) and by (ii) estimating the decrease in genetic signal with key indicators (1), (3), and (4) from Step 1. Please note that key indicators (2) and (5) are not used to evaluate the genetic signal of the latent factor because they are not clearly defined (e.g., heritability is defined as a ratio with phenotypic variance as denominator, which is arguably absent in latent genetic factors).

To evaluate the overall loss of statistical power, we need to make assumptions about the magnitude of the SNP effects. One approach is to compute the squared standardized coefficients^1^, approximated as *r*^2^ = *Z^2^/N*, and then evaluate the median among the subset of genome-wide significant SNPs (*P* < 5×10^−8^) in the down-sampled GWAS. Given that statistical power is the probability of correctly rejecting the null hypothesis when the alternative hypothesis is true, it can be computed as 1-CDF_λ_ [x_1_^2^(c)], where CDF_λ_ is the cumulative distribution function for a χ^2^ distribution with 1 degree of freedom and the non-centrality parameter λ = Nr^2^. The sample size, N, is set to the *EffN* of the summary statistics being evaluated. The term χ^2^(c) is the critical value (∼29.7) at the threshold of genome-wide significance (*P* < 5×10^−8^) for a χ^2^-test with 1 degree of freedom. As a complement, we suggest evaluating the power to detect arbitrary effect-size magnitudes, for which we selected three magnitudes representative of effects reaching genome-wide significance in recent large-scale GWAS (*r*^2^ = 0.003%, 0.004%, or 0.005%).

As in the original study (Karlsson Linnér et al., 2021), we estimated individual SNP effects on the latent EXT-minus-23andMe factor with Genomic SEM, which we refer to as the EXT-minus-23andMe summary statistics. We then evaluated the loss of signal at the factor level (Figure S1–2). We expect the loss of power to be more noticeable at the level of individual loci compared to the follow-up analyses presented, which aggregate genetic signal across larger sets of SNPs or genome wide.

### 4. How similar are gene-property analyses when using down-sampled GWASs?

The biological correspondence of down-sampled univariate or multivariate GWAS can be evaluated by comparing the results from the Multi-marker analysis of genomic annotation (MAGMA) gene-property analyses in the *SNP2GENE* function of Functional Mapping and Annotation of Genome-Wide Association Studies (FUMA; Watanabe et al., 2017); version 1.5.0e) software using Spearman rank correlations of point estimates.

As done in the original paper, we ran gene-property analyses on the EXT minus 23andMe summary statistics to (1) test 54 tissue-specific gene expression profiles, and (2) test gene expression profiles across 11 brain tissues and developmental stages with reference data from BrainSpan (Allen Institute for Brain Science., 2022). We used the default settings of SNP2GENE, which match those used to conduct the gene-based analyses reported in the original study (Karlsson Linnér et al., 2021).

We additionally used FUMA to extract the number of lead SNPs associated with EXT and EXT-minus-23andme. FUMA conducts conventional linkage-disequilibrium (LD) informed pruning (“clumping”) of GWAS summary statistics to count the number of near-independent genome-wide significant lead SNPs. When clumping, FUMA computes LD with the publicly available European subsample of the 1000 Genomes Phase 3 reference panel as the default setting (though, researchers should depart from this default to match the genetic ancestry of the down-sampled GWAS being evaluated). Please note that these analyses differ from the original study in terms of clumping parameters and LD reference panel.

Because power loss is more noticeable at the level of individual SNPs compared to methods that aggregate genetic signal among sets of SNPs or genome-wide, we recommend researchers interested in following up on individual SNPs use the original and not the down-sampled summary statistics for best precision.

### 5. How similar is the pattern of genetic correlations with other traits when using down-sampled GWASs?

To assess the convergent and discriminant validity of down-sampled multivariate GWAS on latent factors, we can examine potential changes in the pattern of genetic correlation with other traits. If the down-sampled analysis tags the same genetic etiology, the confidence intervals of the point estimates should display considerable overlap. The overall pattern can be examined by estimating the rank correlation of the point estimates across traits, whereas significance of changes to individual genetic correlations can be assessed using a *t*-test.

The original study estimated genetic correlations between EXT and 91 other traits (Karlsson Linnér et al., 2021). Here, we performed the same analysis for EXT-minus-23andMe and then examined whether the pattern of genetic overlap was preserved after removing restricted data. Since the summary statistics of some of the 91 traits in the original study include restricted data, we conducted these analyses on the 79 traits with publicly available summary statistics (Table S5).

### 6. How much explanatory power is lost when using polygenic scores (PGSs) constructed from down-sampled GWASs?

Generally, the loss of genetic signal from down-sampling will only exacerbate the problem of measurement error in PGSs constructed with finite-sample estimates as weights (Becker et al., 2021). As one of the most common third-party applications of publicly available GWAS summary statistics, we strongly encourage researchers to evaluate the loss of explanatory power in their main PGS analysis before they share down-sampled summary statistics with other users. This loss can be evaluated (i) across traits, as indicated by the overall reduction in variance explained (*R^2^*/pseudo-*R^2^*) and (ii) with the rank correlation of point estimates to evaluate the comparability of the overall pattern of polygenic score associations.

Following the original study protocol (Karlsson Linnér et al., 2021), we constructed PGSs in two hold-out samples: the Collaborative Study on the Genetics of Alcoholism (COGA; Begleiter, 1995; Bucholz et al., 2017; Edenberg, 2002); N = 7,594) and the National Longitudinal Study of Adolescent to Adult Health (Add Health; Harris et al., 2013; McQueen et al., 2015); N = 5,107). We constructed the PGSs from the EXT-minus-23andMe summary statistics (EXT-minus-23andMe PGS), adjusted for LD with PRS-CS (version 20 October 2019; Ge et al., 2019), which restricts the PGS to ∼1 million HapMap3 SNPs. The default settings are sensible for most standard uses (Bayesian gamma-gamma prior of 1 and .5, and 1,000 Monte Carlo iterations with 500 burn-in iterations).

We compared the explanatory power of the EXT-minus-23andMe PGSs with the one reported in the original study from analyses of a phenotypic externalizing factor, followed by a set of outcomes related to, or affected by, externalizing behaviors and disorders (e.g., smoking initiation, substance-use disorders, or childhood developmental disorders) (Table S6). Linear regression was applied to continuous outcomes and logistic regression to dichotomous outcomes. We evaluated the incremental *R^2^*/pseudo-*R^2^*by subtracting the variance explained by a baseline model with only covariates (age, sex, and the first ten genetic principal components) from the variance explained by a model with the covariates and PGS. Confidence intervals were estimated with the percentile bootstrap method (1,000 iterations). We then evaluated whether the coefficient estimates of the down-sampled EXT-minus-23andMe PGSs were comparable to the estimates of the PGS of EXT from the original paper (Figure 4).

We are aware of recent suggestions to evaluate the squared (semi-)partial correlation in favor of the incremental *R^2^*/pseudo-*R^2^*, but the results of these two alternatives approaches are often highly similar (except when analyzing height). For comparability with the original study, we retained the incremental *R^2^*/pseudo-*R^2^* measure.

## Results

### 1. What is the loss of genetic signal in down-sampled univariate GWASs?

In the initial check of genetic overlap between the full and down-sampled summary statistics of the same trait, we found genetic correlations close to, but still significantly less than unity: 0.966 (SE = 0.007) for SMOK and 0.953 (SE = 0.012) for CANN^2^, which motivated us to apply our approach to evaluate the comparability of the down-sampled summary statistics to those from the original paper.

The loss of genetic signal was evaluated using the five key indicators. First, down-sampling reduced the *EffN* of the two univariate GWASs on SMOK and CANN by about 47% and 12%, respectively (Table 1), which is a marked reduction with potential down-stream consequences. However, down-sampling did not meaningfully impact heritability estimates nor the attenuation/stratification bias ratio, which is important for expecting a comparable factor structure in the multivariate analysis below. Similarly, down-sampling did not meaningfully influence the genetic correlations among the seven indicator phenotypes (Figure 1), which increases the expectation of obtaining a similar factor structure.

Nevertheless, there was a noticeable loss of genetic signal as measured by mean χ^2^ and the genomic inflation factor. The greatest decrease was observed for the down-sampled GWAS on SMOK (Δ mean χ^2^ = 2.06 – 3.15 = –1.09; –34.6%), while the decrease for CANN was less pronounced (–1.3%). Similar decreases were observed for the genomic inflation factor: –25.9% and –1.0% for SMOK and CANN, respectively. The overall stability we observed for the heritability estimates and attenuation ratios suggest that the factor loadings in the down-sampled Genomic SEM factor analysis will resemble those of the original paper (Step 2). The decrease in genetic signal in SMOK and CANN should translate into larger standard errors in the factor analysis and loss of statistical power to detect SNP effects in the multivariate GWAS of EXT-minus-23andMe (Step 3).

### 2. How do the factor loadings and factor model fit differ in multivariate Genomic SEM when the indicator phenotypes are down-sampled univariate GWASs?

The factor loadings, residual variances, and model fit statistics were comparable in the down-sampled single factor solution (Figure 2; Table S2). Neither the factor loadings nor residual variances were statistically different from the original estimates. The largest non-significant difference was observed for the factor loading of the indicator phenotype RISK, which increased from 0.54 (*SE* = 0.03) to 0.56 (*SE* = 0.03). A similar-sized, non-significant decrease was observed for CANN: from 0.77 (*SE* = 0.03) to 0.75 (*SE* = 0.03). Furthermore, the comparative fit index (CFI) and standardized root mean square residual (SRMR) were similar between the down-sampled and original factor models and were within the preregistered thresholds for “good fit” (i.e., CFI > 0.9, and SRMR < 0.08) of the original study (Karlsson Linnér et al., 2021). In our example, we obtain close to identical factor loadings and model fit when applying the best-fitting factor solution of the original study to the empirical genetic covariance matrix of the down-sampled summary statistics.

**Fig 2.**
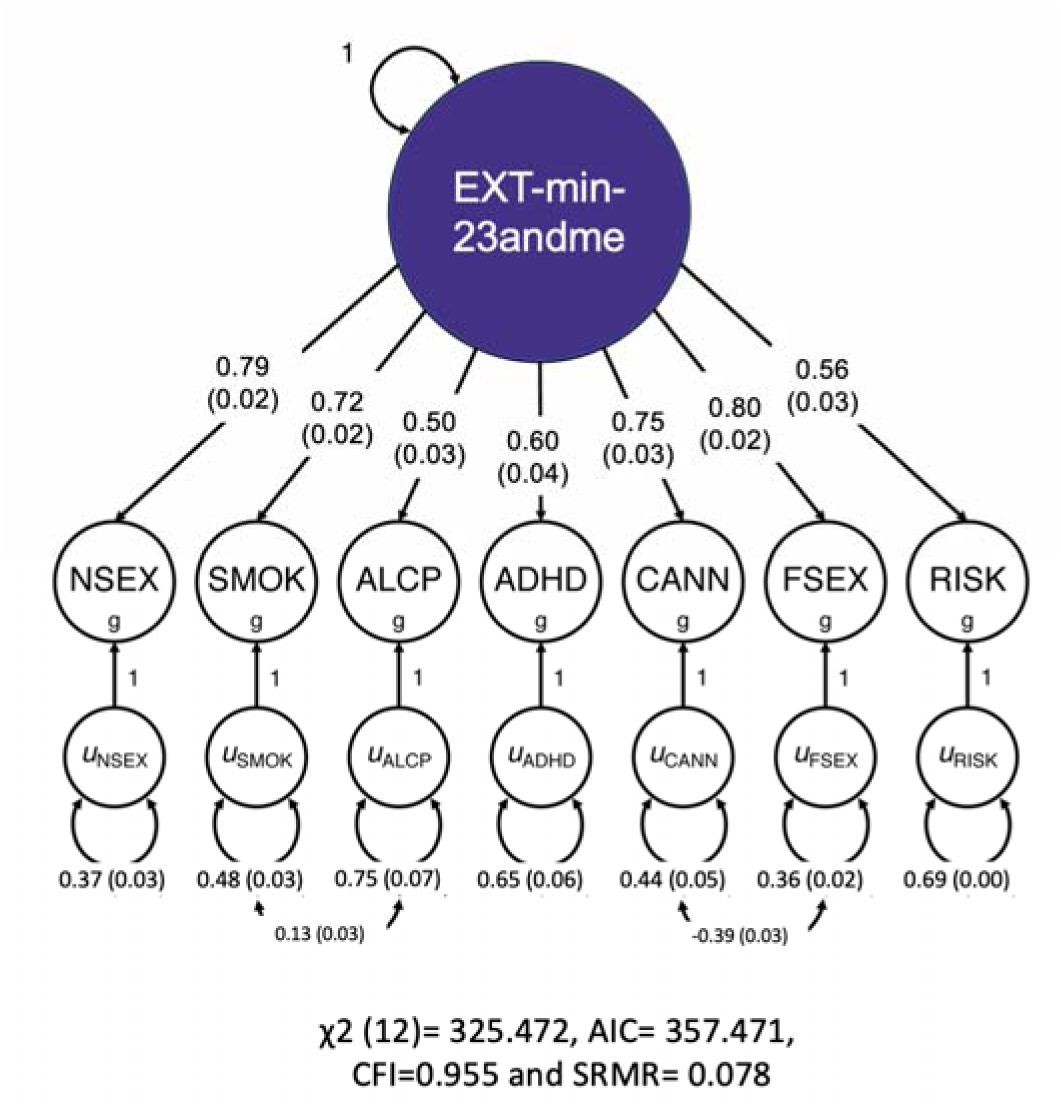
Path diagram of a single-factor model with seven indicator phenotypes, of which SMOK and CANN are down-sampled, as estimated with Genomic SEM. These results are also reported in **Table S2**. The same figure displaying the results of the original study is available here: https://www.nature.com/articles/s41593-021-00908-3/figures/1 Abbreviations: EXT g = genetic externalizing factor; ADHD = attention-deficit/hyperactivity disorder; ALCP = problematic alcohol use; CANN = lifetime cannabis use; FSEX = age at first sexual intercourse (reverse coded); NSEX = number of sexual partners; RISK = risk tolerance; SMOK = lifetime tobacco initiation; AIC = Akaike Information Criterion; CFI = comparative fit index; SRMR = standardized root mean square residual.

### 3. What is the loss of genetic signal at the factor level of multivariate GWAS when the indicator phenotypes are down-sampled univariate GWASs?

We estimated a multivariate GWAS of the EXT-minus-23andMe factor (see Step 2). The genetic correlation between the summary statistics from the multivariate GWAS of EXT and EXT-minus-23andMe was strong but significantly less than unity (*r_g_* = 0.978, *SE* =

0.001), which motivated Steps 4–6. The EffN of the multivariate GWAS of EXT-minus-23andMe was 1,045,957 (about 70.1% of that on EXT). The mean χ^2^s of the EXT and EXT-minus-23andMe factors were 3.12 and 2.37, respectively, corresponding to a 24% decrease. The reduction in the genomic inflation factor was similar (–18%). Thus, there was an appreciable loss of genetic signal in the down-sampled GWAS of EXT-minus-23andMe.

The reduction in mean χ^2^ and genomic inflation factor suggested some loss of power to detect SNP effects. Down-sampling decreased the power by 17.8pp to detect the median of squared standardized coefficients among the genome-wide significant SNPs (i.e., median *r^2^*= 0.0038%), and about 5–45pp less power to detect the three assumed effect-size magnitudes (r^2^ = 0.003%, 0.004%, or 0.005%) (Figures S1–2).

### 4. How similar are the gene-property analyses when using down-sampled GWASs?

We ran gene-property analyses using MAGMA on the EXT-minus-23andMe summary statistics. The Spearman rank correlation of the point estimates from the MAGMA 54 tissues-specific gene expression profiles on the down-sampled and restricted data multivariate GWAS summary statistics was 0.98, suggesting a comparable pattern of gene-tissue expression (Table S3 and Figure S4). The Spearman rank correlation of the point estimates from the MAGMA gene expression profiles across 11 brain tissues and developmental stages also suggested great similarity (*r* = 0.98) (Table S4 and Figure S5). Furthermore, the same 14 tissues, and three developmental stages, remained significant after Bonferroni-correction in the down-sampled analysis (Table S3–4). This evaluation showed that, in the case of EXT-minus-23andMe, the down-sampled gene-property analyses led to similar biological insights as those from the original paper (Karlsson Linnér et al., 2021).

Pruning of the summary statistics to near-independent lead SNPs (using the FUMA default settings), identified 358 lead SNPs for EXT-minus-23andMe, as compared to 825 lead SNPs for EXT. Note that the number of lead SNPs reported here for EXT differs from the original study because that study used a restricted-access genetic reference panel and different settings for the pruning parameters. In our scenario, down-sampling reduced the number of near-independent lead SNPs by 56.6%. Therefore, we recommend that users interested in following up on individual genome-wide significant SNPs associated with externalizing prioritize the version with 23andMe data.

### 5. How similar is the pattern of genetic correlations with other traits when using down-sampled GWASs?

We assessed the pattern of genetic correlations of EXT-minus-23andMe with other traits and found this pattern to nearly identical to that of the original study (Spearman *r* ∼ 1) (Figure 3, Table S5). Furthermore, none of the point estimates were statistically different. Thus, in our scenario, down-sampling did not meaningfully impact the genetic correlations with other traits, meaning that researchers interested in such analyses can safely proceed with using the down-sampled summary statistics.

**Fig 3.**
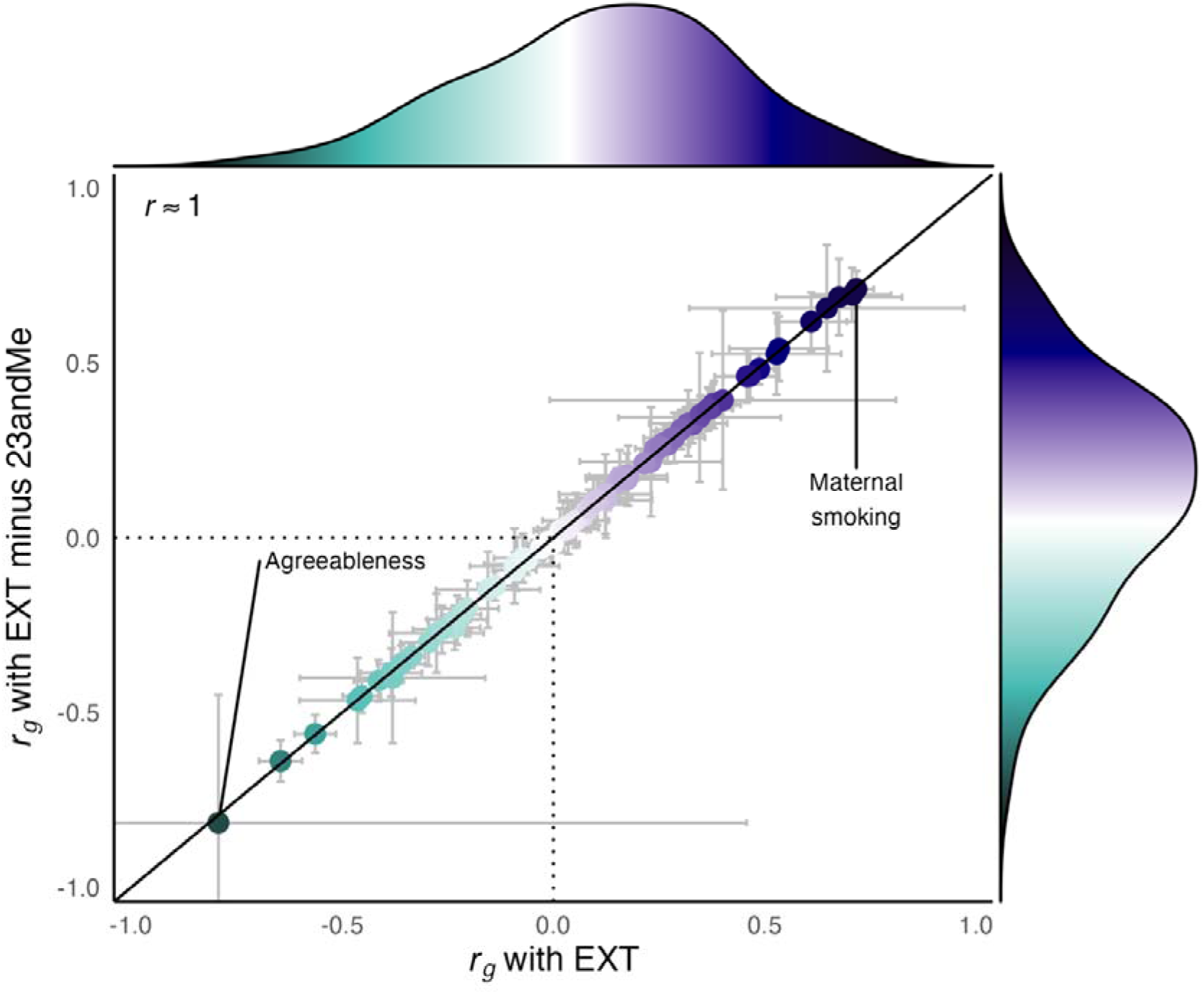
Scatterplot of genetic correlations (*r_g_*) and marginal density plots between EXT (y-axis) or EXT-minus-23andMe (x-axis) with 77 other phenotypes. Each point corresponds to the genetic correlation coefficient with its 95% confidence intervals (*r_g_* + 1.96 × *SE*) estimated with bivariate LD Score regression. **Table S5** reports the estimates, their standard errors, and confidence intervals. The Spearman rank correlation reported in the figure is rounded from *r* = 0.9995. No particular shape, such as a normal distribution, is expected for the marginal density because the figure displays an arbitrary selection of traits.

### 6. How much explanatory power is lost when using polygenic scores (PGSs) constructed from down-sampled GWASs?

The down-sampled PGS for EXT-minus-23andMe explained 8.4% and 8.5% of the variance of a phenotypic externalizing factor in Add Health and COGA, respectively, which is 1.9pp and 0.5pp less compared to the same analysis in the original study (Table S6). The overall reduction in explanatory power across other outcomes was less pronounced, on average 0.35pp in Add Health, and 0.23pp in COGA. The largest decrease was observed for lifetime smoking initiation with 2.1pp and 1.7pp, followed by lifetime cannabis use with 1.1pp in Add Health (but only 0.55pp in COGA), which may be explained by these two indicator phenotypes being most affected by the down-sampling. For most other traits, the variance explained by the down-sampled PGS was comparable to the original study.

Secondly, the Spearman rank correlation of the regression coefficients was 0.996, suggesting great similarity in point estimates (Figure 4). All the coefficients of the down-sampled PGS fell within the confidence intervals of their original study counterparts (Table S6), except those for the phenotypic externalizing factor (in Add Health), lifetime smoking initiation, and lifetime cannabis use (in Add Health). Overall, our down-sampled polygenic score results were comparable to those from the original study, meaning that researchers interested in using the down-sampled summary statistics to construct PGS for EXT-minus-23andMe can generally expect similar results. However, we recommend the users be aware of the weaker explanatory power for certain outcomes.

**Fig 4.**
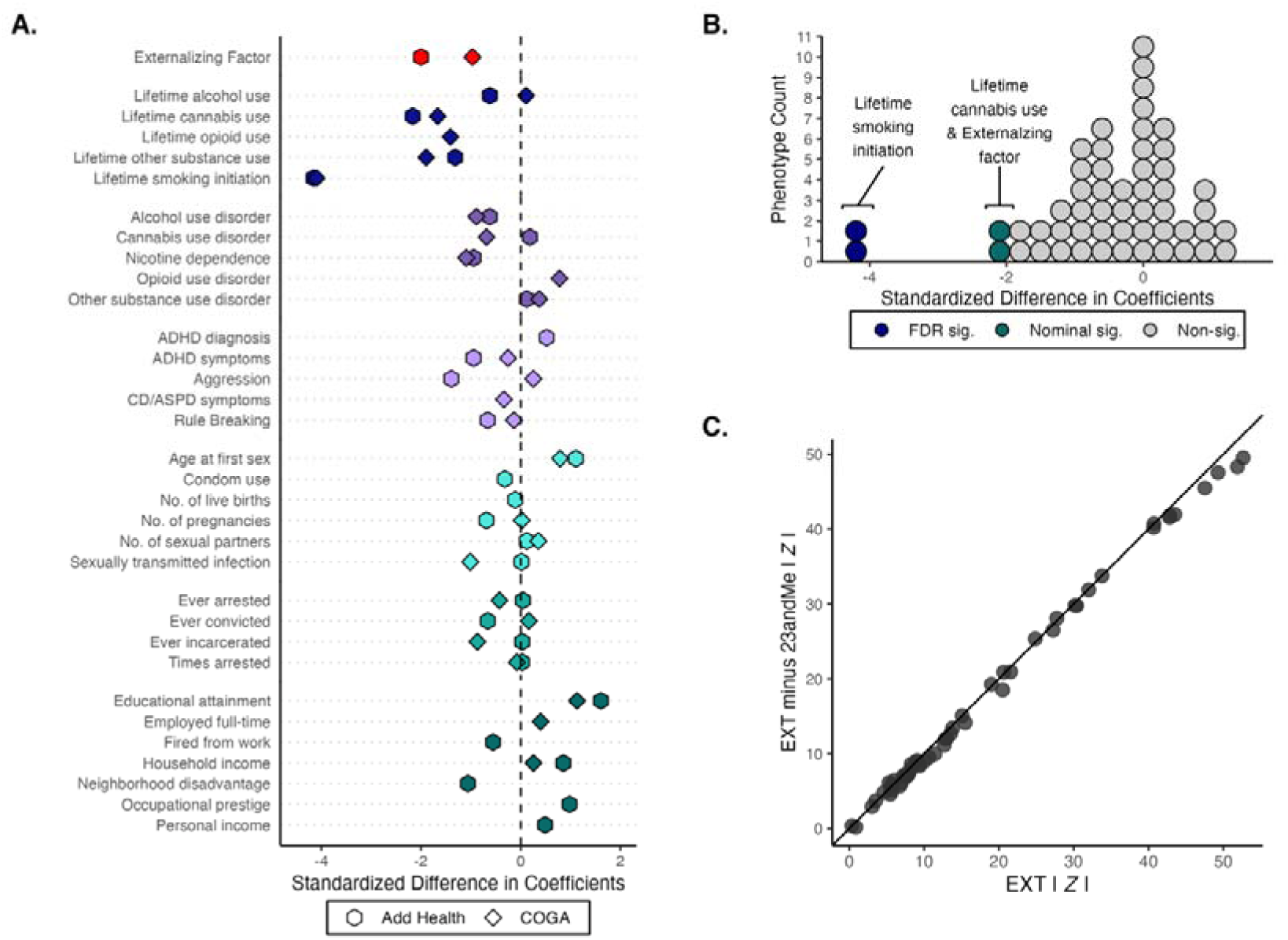
Comparison of the down-sampled polygenic score (PGS) analyses in Add Health (29 phenotypes) and the Collaborative Study on the Genetics of Alcoholism (COGA; 26 phenotypes). Panel A displays the standardized difference between the coefficient estimates (i.e., a *Z*-statistic) of the down-sampled PGS for EXT-minus-23andMe versus the PGS for EXT from the original study. Absolute values were evaluated so that a negative standardized difference refers to an attenuation towards zero in the down-sampled analysis. Panel B displays the same measure but as a histogram. Four coefficient estimates were significantly (at the 5% level) attenuated in the down-sampled analysis: lifetime smoking initiation (Add Health and COGA; *P* = 3.18×10^−5^ and 4.17×10^−5^, respectively), the phenotypic externalizing factor (Add Health; *P* = 0.046), and lifetime cannabis use (Add Health, *P* = 0.03). None of the coefficients were significantly larger in the down-sampled analysis. Panel C displays a scatter plot of the absolute value of the coefficient estimates divided by their respective standard errors (i.e., a *Z*-statistic). These results are also reported in **Table S6**.

## Discussion

Unrestricted access to data and results is the cardinal tenet of open science. Here, we propose a systematic approach (i) to evaluate the comparability of down-sampled GWAS summary statistics with their restricted data counterparts, and (ii) to assess the impact of using down-sampled univariate summary statistics in multivariate GWAS with Genomic SEM. We examined the loss of genetic signal in down-sampled univariate GWAS (Step 1), the change in the factor model loadings and fit (Step 2), the loss of genetic signal at the factor-level of down-sampled multivariate GWAS (Step 3); and for potential changes to gene-property analyses (Step 4), the pattern of genetic correlations with other traits (Step 5), and the explanatory power of polygenic score analyses in independent samples (Step 6).

We applied these steps to the largest available multivariate GWAS of externalizing to evaluate the quality and predictive performance of the results following restricted data removal. We found nearly identical model fit and parameter estimates, genetic correlations with other phenotypes, and polygenic score analyses of externalizing phenotypes in independent samples. As expected, we observed a decrease in power and genetic signal in the down-sampled univariate and multivariate summary statistics. Although fewer lead SNPs were identified for EXT-minus-23andMe compared to EXT, the genes associated with EXT and EXT-minus-23andMe were similar in terms of region and developmental timing of expression. In the PGS context, EXT and EXT-minus-23andMe performed similarly well. Therefore, while we suggest that the down-sampled summary statistics may be used in analyses related to gene enrichment, genetic correlations, or polygenic scores, the summary statistics with restricted data should be prioritized for gene identification or following up on genome-wide significant hits.

In our example, removing restricted data did not change the construct that was identified by genetic factor analysis: The genetic correlation between the factor identified without 23andMe data and the factor identified with 23andMe data was near unity, and the factors had highly similar associations with external variables. But this outcome is not guaranteed. Removing restricted data may be more impactful for univariate GWASs prior to their inclusion in meta-analyses and multivariate GWAS with different indicator phenotypes and model structures. The consistency we observed between EXT and EXT-minus-23andMe is likely explained by the inclusion of restricted data in only a subset of indicators, with just one of seven summary statistics experiencing a substantive reduction in genetic signal (i.e., 35% decrease in the mean χ^2^ of SMOK). In the circumstance that more indicators had included 23andMe data, we could have expected greater discrepancies between EXT and EXT-minus-23andMe.

The issues raised here are also relevant in the context of GWAS meta-analyses. Removing a restricted set of cohort-level summary statistics from a single-phenotype GWAS meta-analysis should mainly affect power if the genetic correlation between the cohort-level summary statistics is close to unity. However, considering that genetic correlations between cohort-level GWASs of the same trait can be substantially less than unity (Levey et al., 2021), removing a large cohort from the meta-analysis can change the genetic etiology of the trait being studied (de Vlaming et al., 2017). Researchers should thus use the approach presented here to examine potential changes in a phenotype’s genetic etiology alongside the expected power reduction after removing a sample from their GWAS meta-analysis. To our knowledge, this has only been done by one meta-analysis (Coleman et al., 2020), where the authors conducted a subset of the steps described in the present study (e.g., changes in heritability, genetic correlations with external variables, and gene enrichment analyses). Therefore, the utility of our systematic approach goes beyond the Genomic SEM context, as some of these steps may apply to other multivariate GWAS implementations.

Providing public summary statistics to the wider research community is crucial to facilitating open science and advancing behavioral and biomedical research. The first step in this process should be to evaluate the comparability of down-sampled summary statistics and their restricted data counterparts. Herein, we provide a systematic approach to investigators who resort to sharing down-sampled GWAS summary statistics and recommend they report these analyses as accompanying documentation to facilitate open science and data sharing.

## Additional information and acknowledgements

### Statement of Ethics

● This study included only secondary data analysis of de-identified data and was not subject to an institutional review board (IRB) review.
● All participants provided written informed consent in the original studies from which these data were drawn. In addition, data collection of each cohort was approved by a review board at each respective institution.

### Conflict of Interest Statement

No competing interests declared

## Acknowledgments & Funding Sources

This research was conducted by the Externalizing Consortium. The Externalizing Consortium has been supported by the National Institute on Alcohol Abuse and Alcoholism (R01AA015416 – administrative supplement to DMD), and the National Institute on Drug Abuse (R01DA050721 to DMD). Additional funding for investigator effort has been provided by K02AA018755, U10AA008401, P50AA022537 to DMD, R01AA029688, and 28IR-0070 to AAP and **T29KT0526 and T32IR5226 to NCK** and SSR from the Tobacco-Related Disease Research Program (TRDRP), NIDA DP1DA054394 to SSR, **R25MH081482-16 to NCK**, R01HD092548 to KPH, as well as a European Research Council Consolidator Grant (647648 EdGe) to PDK. The content is solely the responsibility of the authors and does not necessarily represent the official views of the above funding bodies. The Externalizing Consortium would like to thank the following groups for making the research possible: 23andMe, Add Health, Vanderbilt University Medical Center’s BioVU, Collaborative Study on the Genetics of Alcoholism (COGA), the Psychiatric Genomics Consortium’s Substance Use Disorders working group, UK10K Consortium, UK Biobank, and Philadelphia Neurodevelopmental Cohort.

## Data Availability Statement

The code for EXT-minus-23andMe is available on the wiki (https://github.com/Camzcamz/EXTminus23andMe/wiki) and the EXT-minus-23andMe summary statistics are available on the externalizing website (https://externalizing.rutgers.edu/ext-23andme-summary-statistics-now-available/).

---

**Figure S1.**
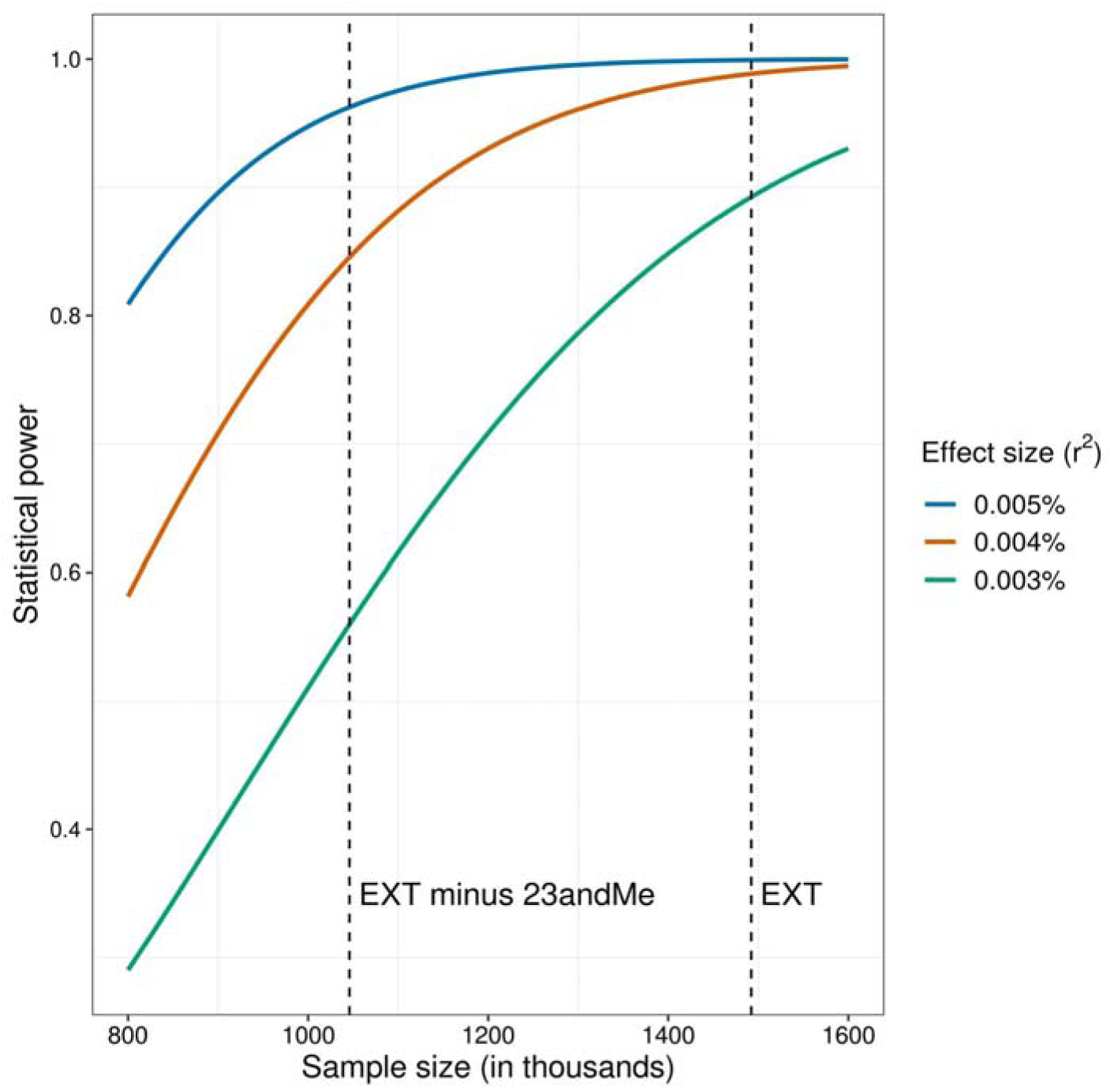
Power analysis of down-sampled GWAS summary statistics. The figure displays the statistical power to detect three arbitrary effect-size magnitudes at genome-wide significance (*P* < 5×10^−8^) as a function of sample size. The three magnitudes were selected to represent smaller magnitudes that reach genome-wide significance in recent large-scale GWAS. The dashed lines mark the sample size of the original multivariate GWAS of EXT (*EffN* = 1,492,085), and that of the down-sampled multivariate GWAS of EXT-min-23andMe (*EffN* = 1,045,957).

**Figure S2.**
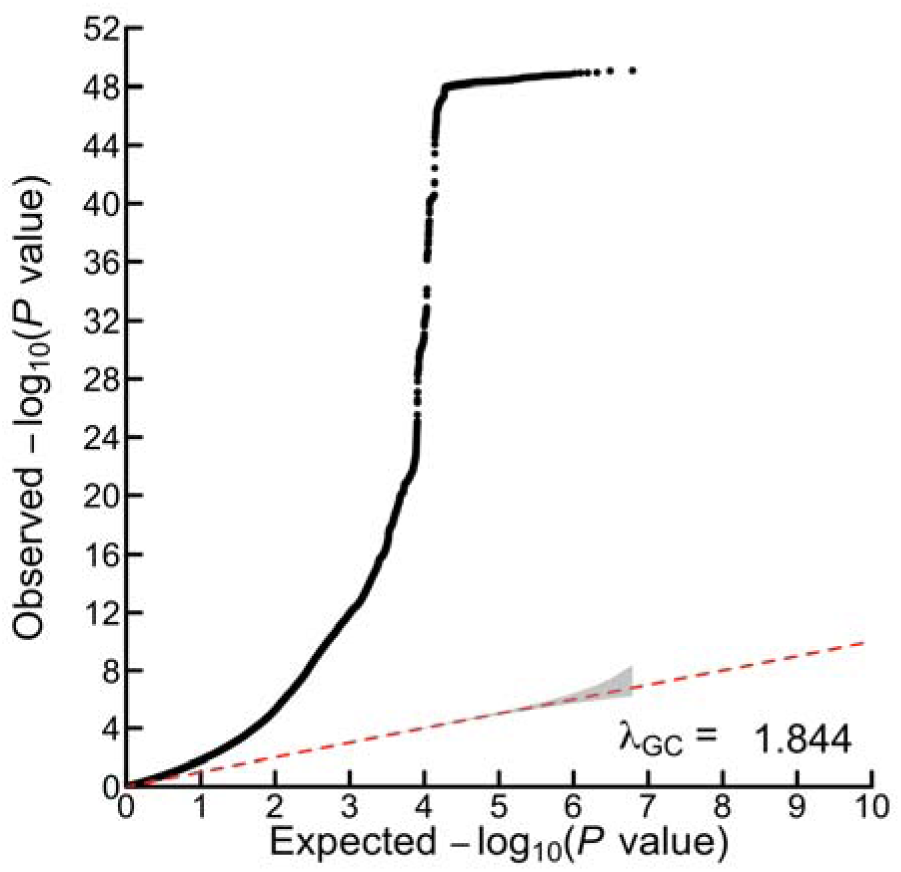
Quantile-quantile (Q-Q) plot. The y-axis shows the observed association *P* value on the –log10 scale (for a two-sided *Z*-test) among the 6,170,305 SNPs in the multivariate GWAS of EXT-min-23andMe (*EffN* = 1,045,957), which are plotted against the expected –log10(*P*) of the null distribution. The gray shaded area shows 95% confidence intervals centered on the null distribution. The genomic inflation factor in the figure, λ_GC_, is the median χ^2^ association test statistic divided by the expected median of the χ^2^ distribution with 1 degree of freedom. This estimate of λ_GC_ differs somewhat from that of LD Score regression, which estimates this statistic using only ∼1 million SNPs. The same plot for the original study is available here: https://www.nature.com/articles/s41593-021-00908-3/figures/6

**Figure S3.**
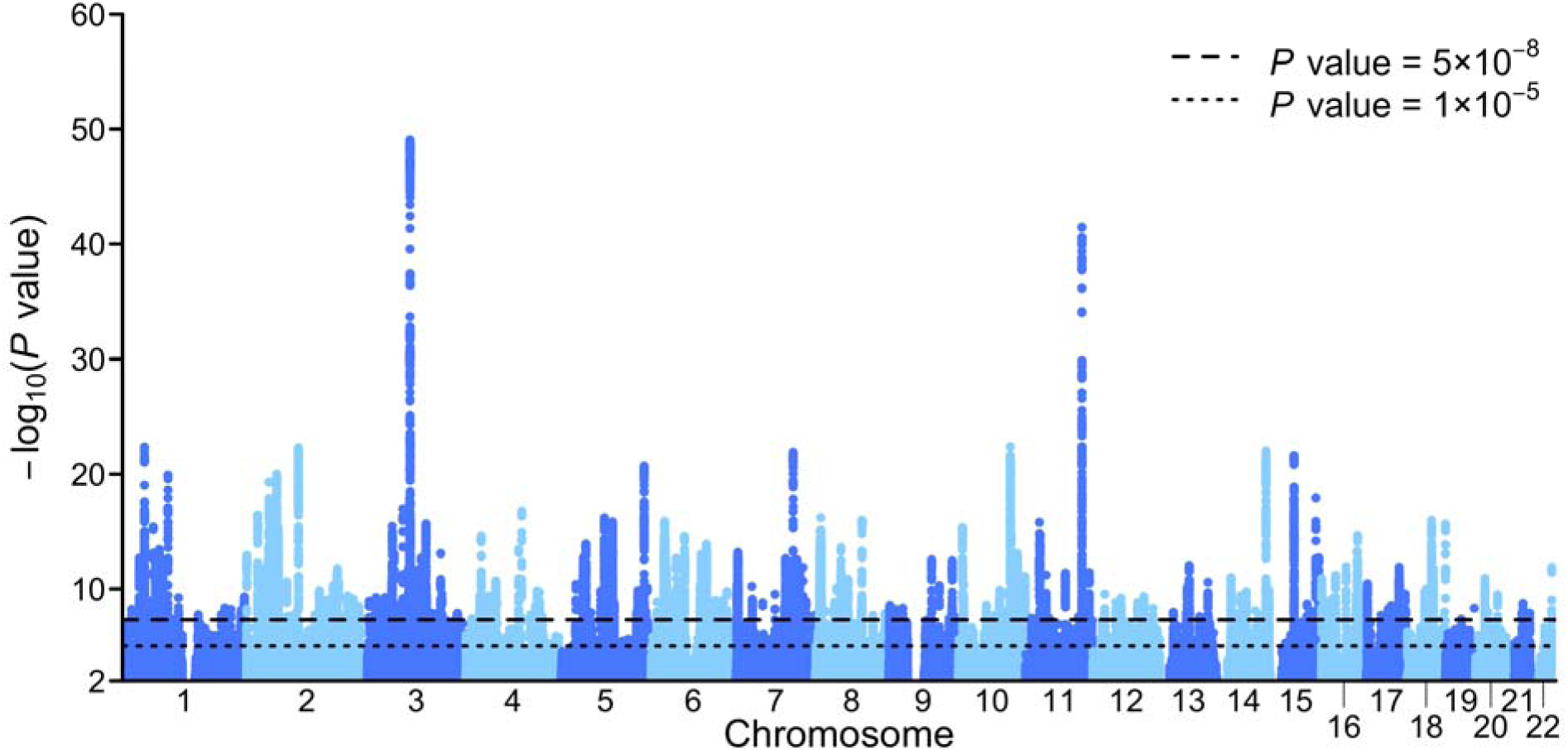
Manhattan plot of the GWAS of the EXT-min-23andMe factor, estimated with Genomic SEM (*EffN* = 1,045,957). The figure displays association *P* values on the –log10 scale (two-sided) for 505,254 with *P* < 0.01 out of 6,170,304 SNPs tested for association. The dashed line represents genome-wide significance (*P* < 5×10^−8^) and the dotted line shows suggestive significance (*P* < 1×10^−5^). A Manhattan plot of the corresponding GWAS analysis in the original study of EXT is available here: https://www.nature.com/articles/s41593-021-00908-3/figures/2

**Figures S4.**
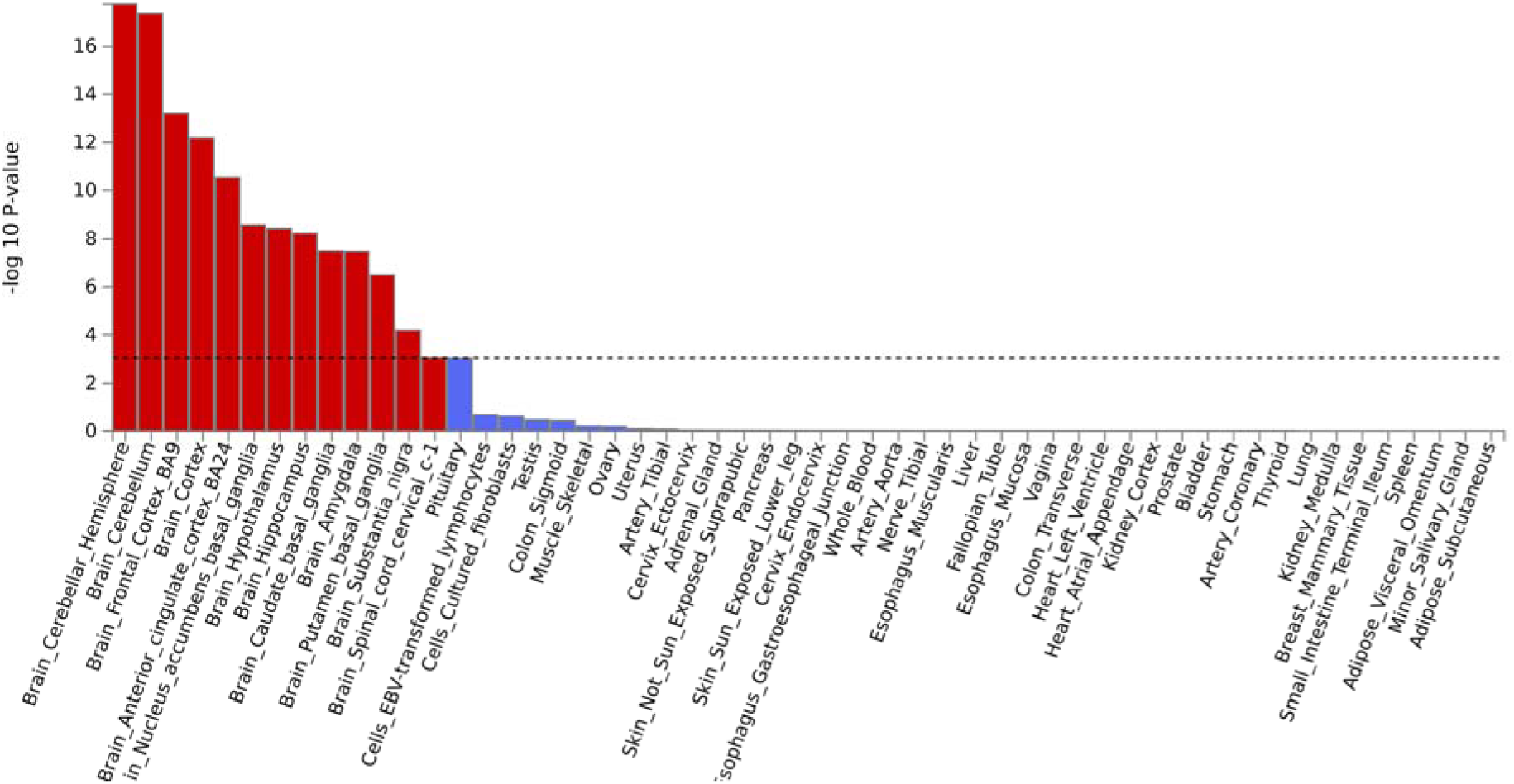
Bar plot of MAGMA gene-property analysis of enrichment in 54 bodily tissues. The figure displays *P* values on the –log10 scale (from one-sided *Z*-tests) of the point estimate from a generalized least squares regression, estimated with MAGMA as implemented in FUMA. The analysis was applied to the summary statistics from the down-sampled multivariate GWAS of EXT-min-23andMe (*EffN* = 1,045,957). Dashed line denotes Bonferroni-corrected significance, adjusted for testing 54 tissues (one-sided *P*<9.26×10^−4^). These results are also reported in **ST3**. The same 14 tissues identified in the original study were also found significantly associated with the down-sampled multivariate GWAS of EXT-min-23andMe. The same plot for the original study is available here: https://www.nature.com/articles/s41593-021-00908-3/figures/9

**Figure S5.**
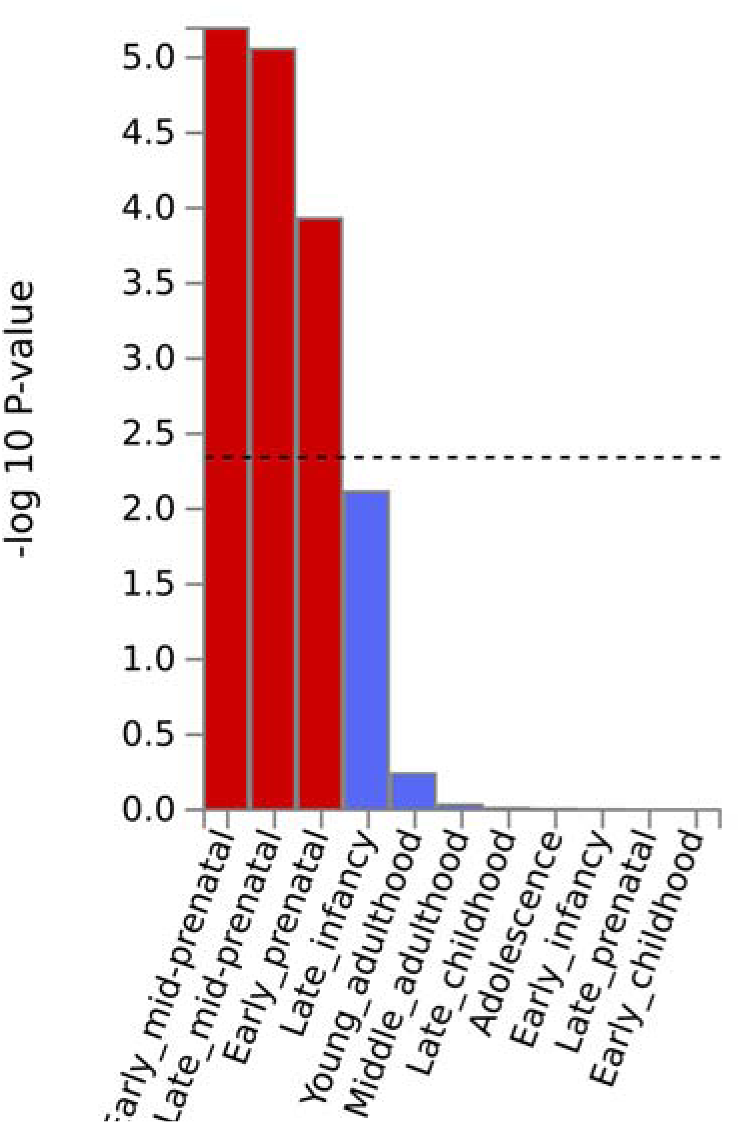
Bar plot of MAGMA gene-property analysis of enrichment in brain tissues across 11 developmental stages (BrainSpan). The figure displays *P* values on the –log10 scale (from one-sided *Z*-tests) of the point estimate from a generalized least squares regression, estimated with MAGMA as implemented in FUMA. The analysis was applied to the summary statistics from the down-sampled multivariate GWAS of EXT-min-23andMe. These results are also reported in **ST4**. Dashed line denotes Bonferroni-corrected significance, adjusted for testing 11 developmental stages (one-sided *P*<4.55×10^−3^). The same three developmental stages identified in the original study were also found significantly associated with the down-sampled multivariate GWAS of EXT-min-23andMe. The same plot for the original study is available here: https://www.nature.com/articles/s41593-021-00908-3/figures/10

## Footnotes

1 An approximate measure of variance explained (R^2^), standardized with respect to the outcome.

2 Estimated with the chi-square cut-off set to 30, i.e., the default cut-off applied by bivariate LD Score regression when estimating the heritability. To our knowledge, there is no consensus on the best cut-off to use.

## Notes

https://github.com/Camzcamz/EXTminus23andMe

### Competing Interest Statement

The authors have declared no competing interest.

